# Sequencing Chemically Induced Mutations in the Mutamouse Lacz Reporter Gene Identifies Human Cancer Mutational Signatures

**DOI:** 10.1101/858159

**Authors:** Marc A. Beal, Matt J. Meier, Danielle LeBlanc, Clotilde Maurice, Jason O’Brien, Carole L. Yauk, Francesco Marchetti

## Abstract

Transgenic rodent (TGR) models use bacterial reporter genes to quantify *in vivo* mutagenesis. Pairing TGR assays with next-generation sequencing (NGS) enables comprehensive mutation spectrum analysis to inform mutational mechanisms. We used this approach to identify 2,751 independent *lacZ* mutations in the bone marrow of MutaMouse animals exposed to four chemical mutagens: benzo[a]pyrene, *N*-ethyl-*N*-nitrosourea, procarbazine, and triethylenemelamine. We also collected published data for 706 *lacZ* mutations from eight additional environmental mutagens. We demonstrate that *lacZ* gene sequencing generates chemical-specific mutation signatures observed in human cancers with established environmental causes. For example, the mutation signature of benzo[a]pyrene, a potent carcinogen in tobacco smoke, matched the signature associated with tobacco-induced lung cancers. Our results show that the analysis of chemically induced mutations in the *lacZ* gene shortly after exposure provides an effective approach to characterize human-relevant mechanisms of carcinogenesis and identify novel environmental causes of mutation signatures observed in human cancers.

## INTRODUCTION

Transgenic rodent (TGR) mutation reporter models have enabled unprecedented insights into spontaneous and chemically induced mutagenesis^1^. Studies of over 200 chemicals, including more than 90 carcinogens, have demonstrated that TGR models offer high sensitivity and specificity for identifying mutagenic carcinogens^1,2^. One of the most commonly used TGR models is the MutaMouse whose genome was recently sequenced^3^. The MutaMouse harbors ~29 copies of the bacterial *lacZ* transgene on each copy of chromosome 3^4^. This is a neutral, transcriptionally-inert reporter gene carried on a shuttle vector that can be recovered from any cell type and transfected into a bacterial host to detect somatic or germline mutations that occurred *in vivo*^5^,^6^. A major advantage of TGR models is the possibility to sequence mutants in order to characterize mutation spectra. This information is necessary to understand mutational mechanisms associated with mutagen exposure and response in different tissues, life stages genetic backgrounds or other contexts. Advances in next-generation sequencing (NGS) technologies have enabled rapid and accurate characterization of TGR mutants^7^,^8^, and integrated TGR-NGS approaches have been used to sequence thousands of mutations^8^,^9^ at a fraction of the cost of whole genome sequencing. Thus, TGR-NGS approaches currently provide a unique methodology for simultaneously assessing the magnitude of the mutagenic response and mutation spectrum to inform underlying mechanisms.

Somatic mutation analysis by NGS has greatly advanced our understanding of the mutational processes operating in human cancers. Algorithms have been developed to mine the extensive database of single nucleotide variations (SNVs) in cancer genomes to identify mutational signatures contributing to individual cancers^10,11,12^. These signatures represent a computationally derived prediction of the relative frequencies of mutation types induced by processes that contribute to all observed mutations within The Cancer Genome Atlas datasets (TCGA; https://www.cancer.gov/about-nci/organization/ccg/research/structural-genomics/tcga). As opposed to standard mutation spectrum characterization that simply describes the frequency of individual nucleotide changes, mutational signatures incorporate flanking nucleotide context. Originally, 30 mutational signatures from 40 different cancer types were identified and reported in the Catalogue of Somatic Mutations in Cancer (COSMIC) database^13,14^. This database was recently expanded to include 71 cancer types and 77 signatures, including 49 single base substitution (SBS) signatures, 11 doublet base substitution (DBS) signatures, and 17 small insertion and deletion (ID) signatures^15^. Each signature encompasses 96 possible mutation types (i.e., 6 possible base pair alterations × 4 different 5’ bases × 4 different 3’ bases). Many of these signatures have been attributed to endogenous processes, but chemical mutagens also play a major contributing role in certain signatures^16^. For example, SBS 4 signature is observed in lung cancer and has been attributed to tobacco smoke^16,17^. This signature has been recapitulated by exposing murine embryo fibroblasts to benzo[a]pyrene (BaP)^18,19^, a major mutagenic component of tobacco smoke. However, several of the mutational signatures currently have no known endogenous or exogenous causative agents^17^; thus, identification of exogenous environmental exposures that contribute to these mutational signatures may aid in elucidating carcinogenic mechanisms.

The pattern of mutations observed in a fully developed cancer is a composite of the signature of the molecular initiating events in the early stages of tumour formation and signatures arising as a result of genomic instability in the evolving tumour^20^. For example, a tumour that originates in the lung of a smoker will have a mutational fingerprint that is caused primarily by DNA damage induced by the many mutagenic compounds found in tobacco smoke^21^. In addition, the person’s age at the time of tumour formation will also determine the contribution of “clock-like” signatures, caused by lifetime DNA replication, to the fingerprint of the tumour^22^. There is now compelling evidence that analysis of the spectrum of mutations in a cancer can provide clues to past environmental exposures that contributed to the development of the cancer^23,24^. Implicit in this is that the exposure signature should be present in the normal tissue before the carcinogenic process becomes apparent. Indeed, previous studies have demonstrated that mutational signatures observed in aflatoxin-induced cancers are observed in normal tissues long before tumour formation^25, 26^. Recent work *in vivo*^27^ and *in vitro*^28^ has shown that chemical-specific signatures detected shortly after exposure match signatures seen in human cancers. Thus, characterization of short-term mutational signatures in non-tumour tissues is a valuable approach to elucidate human-relevant mechanisms of carcinogenesis.

In this study, we used TGR-NGS to characterize mutations induced by four established mutagens to determine if these mutation profiles inform carcinogenic mechanisms within COSMIC signatures. For this purpose, we chose four chemicals with varying mutagenic potencies, mode of action, and carcinogenic classification (as determined by the International Agency for Research on Cancer): one known class 1 carcinogen, BaP; two probable class 2 carcinogens including *N*-ethyl-*N*-nitrosourea (ENU) and procarbazine (PRC); and one class 3 chemical with inadequate information to be classified, triethylenemelamine (TEM). MutaMouse males were exposed by gavage to the chemicals or solvent for 28 days and DNA was collected from bone marrow for analysis. To further compare *lacZ* mutation spectra and COSMIC signatures, published Sanger sequencing data from 17 studies involving eight mutagens were also examined (Supplementary Table S1). These studies include data from mice exposed to electromagnetic radiation^29,30,31,32,33^, alkylating agents and adduct-forming agents^34,35,36,37,38,39,40^, and a nitrogenous base analog^41^. Data from control animals in these studies and others^42,43,44,45,46^ were also included to generate a background mutation signature. Using *lacZ*-derived mutation data, we validated COSMIC signatures with proposed aetiologies through the identification of the expected signatures in the relevant exposure groups. We argue that analysis of COSMIC signatures observed in exposed animals can be used to generate or test hypotheses of mutagenic mechanisms associated with human mutational signatures of unknown etiology.

## RESULTS

We used mutation spectra generated in-house for four chemicals (BaP, ENU, PRC, and TEM) and vehicle matched controls, and published data from eight agents, including BaP and ENU (Supplementary Table S1) and their matched controls, to query the COSMIC database and elucidate the role of environmental mutagens in cancer development. The overall experimental design is summarized in Figure 1.

**Figure 1.**
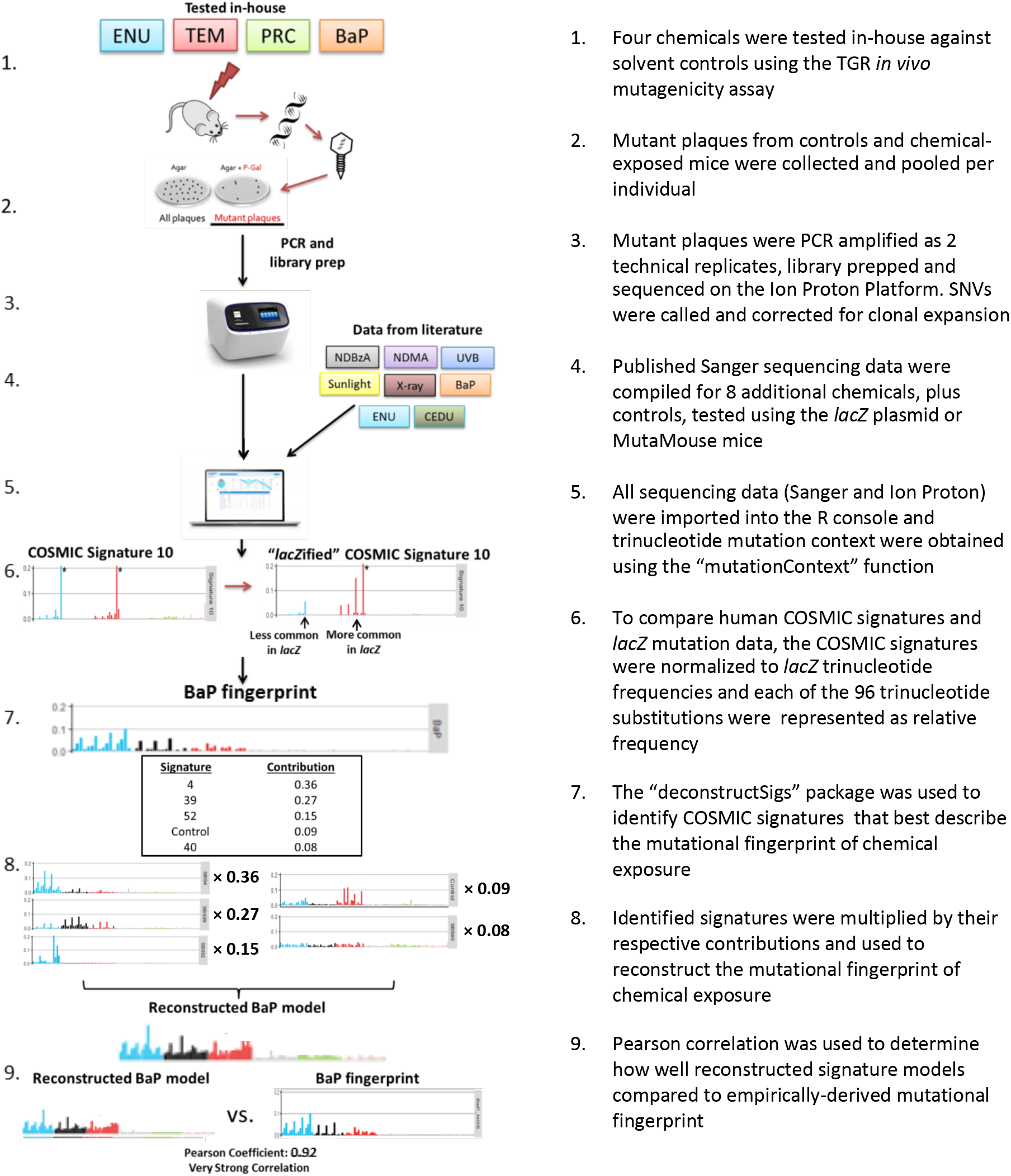
Experimental design. The experimental workflow included: animal exposure and determination of mutant frequencies (Steps 1-2); sequencing of collected plaques and collection of published *lacZ* sequenced data (Steps 3-4); generation of mutation profiles (steps 5-6); and query of the COSMIC database to identify mutational signatures that contributed to the mutation profile of tested agents (Steps 7-9).

Mutation spectra were generated from plaques collected during experiments aimed at evaluating the induction of mutations in the bone marrow of MutaMouse males exposed to either BaP, ENU, PRC, or TEM using the *lacZ* assay^5^. Mutant frequencies were previously reported for BaP^8^, PRC^47^ and TEM^48^, while mutant frequencies are reported here for the first time for ENU. All of the exposures caused increases in mutant frequencies relative to vehicle-matched controls (Supplementary Table 2), and the results were highly significant (P < 0.0001) for BaP (122.9-fold), PRC (9.7-fold), and ENU (7.2-fold). TEM exposure also increased mutant frequency relative to controls (1.6-fold; P = 0.048), but it was less potent than the other agents. The potency ranking of exposures (BaP > PRC/ENU > TEM) was consistent with expectations.

### Mutation Characterization and Spectral Analysis

Sequencing of 5,419 mutant plaques from bone marrow DNA enabled the characterization of 2,751 independent mutations (Supplementary Table S3). Sequenced plaques from BaP, ENU, and controls were generated by both NGS and Sanger sequencing. Specifically, there were 1105, 406, and 438 mutations identified by NGS for BaP, ENU, and Controls, respectively. The corresponding numbers were 60, 207, and 508 for Sanger sequencing. The mutation spectra generated by the two sequencing approaches were consistent for each of the three groups (data not shown). Thus, within each group, the two sets of mutations were combined. Overall, there were 1,046, 2,914, 129, 902, and 428 mutants sequenced in the Controls, BaP, PRC, ENU, and TEM groups, respectively. These sequenced mutants represented 512, 1,547, 120, 419, and 153 independent mutations in the five groups, respectively.

In the *lacZ* gene, there are 3,096 positions × 3 possible substitutions at each position for a total of 9,288 possible unique SNV events; however, not all of these can be detected using a functional assay, since many result in silent mutations. Sequencing mutants from the different groups identified 891 unique SNVs, 338 of which overlapped between two or more groups (Supplementary Figure S1). Specific to each group, there were 55, 377, 14, 85, and 22 unique SNVs for Controls, BaP, PRC, ENU, and TEM, respectively (Supplementary Table S3). The mutations detected in this study are limited almost exclusively to point mutations and small indels (1-21 bp), as large deletions are infrequently recovered during packaging of the DNA for the *lacZ* assay^8^.

The mutation spectra of the four chemicals were significantly different from the control mutation spectrum (Figure 2; P ≤ 0.0008). The COSMIC convention is to represent mutations based on pyrimidine changes; thus, we present our mutation spectrum using the same convention. The main spontaneous mutation is represented by C>T transitions, which are thought to arise through spontaneous mechanisms such as deamination of methylated cytosines^49^. Although there may be proportional declines in specific mutations relative to controls (Figure 2), all of the chemicals tested in this study, with the exception of TEM, increased the mutation frequency of substitutions (e.g., C>T; Supplementary Figure S2).

**Figure 2.**
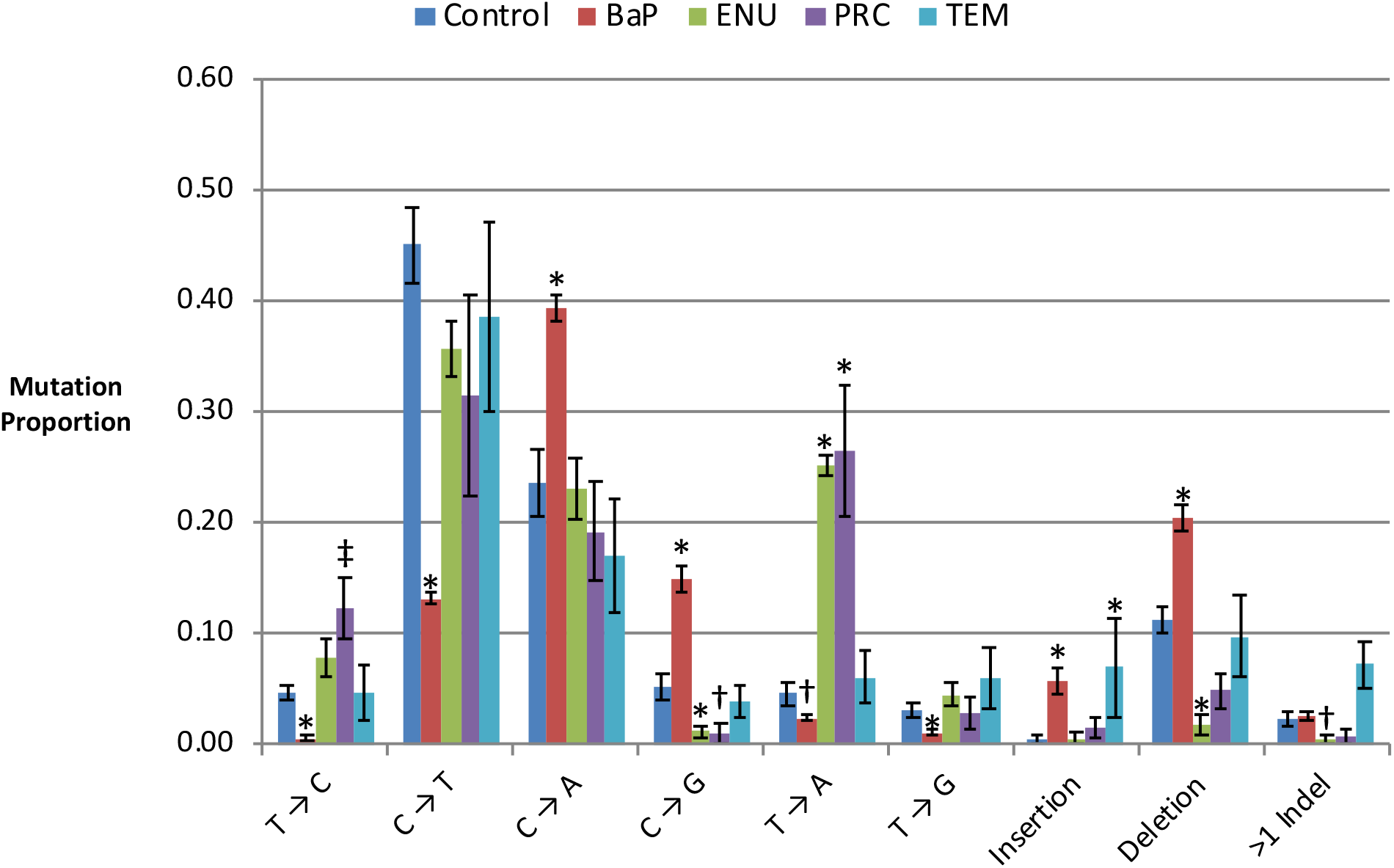
Spontaneous and chemical-induced mutation proportions in bone marrow as characterized by NGS. BaP, shown in red has significantly higher proportions of C>A, C>G, insertions, and deletions compared to control. In contrast, there is a lower proportion of T>C, C>T, T>A, and T>G mutations than control. ENU, shown in green, has a higher proportion of T>A mutations, while C>T, C>G, and deletions are lower. PRC, shown in purple, has a higher proportion of T>A compared to control, and a marginally significant increase in T>C mutations compared to control (P = 0.055). The mutation spectrum for TEM, shown in turquoise, is most similar to that of the control, with the exception of a significant increase in the proportion of insertions. ‡ P < 0.1, † P < 0.05, * P < 0.0001.

The mutation spectra of BaP and ENU are consistent with previous observations. BaP exposure caused cytosine transversions and indels (Figure 2), mainly C>A SNVs, consistent with the formation of bulky DNA adducts mostly at the N2 of guanine^8^. ENU induced T>A mutations consistent with alkylation of thymine, specifically O^2^-and O^4^-ethyl thymine^50, 51^. We found that PRC induced T>A mutations and, to a lesser extent T>C mutations, which is consistent with the pattern of mutations that was observed in an endogenous gene^52^. The mutation spectrum of TEM was significantly different from controls, but there were no significant changes in specific SNV types. Instead, this effect is mainly driven by the higher proportion of TEM-induced single nucleotide insertions compared to control animals. TEM also induced the highest proportion of >1bp indels among all chemicals tested (Figure 2).

### Identification of COSMIC Signatures Using *lacZ* Mutations

We explored the use of the *lacZ* sequence to obtain mutational signatures associated with human cancers. Although the COSMIC database (version 3) includes also DBS and ID signatures, we focused on SBS signatures because the *lacZ* assay detects almost exclusively these types of events. We first divided each trinucleotide frequency in the *lacZ* transgene (Figure 3) by the respective human genome frequencies (hg38) to create a *lacZ*-normalized set of the 49 COSMIC SBS signatures (Supplementary Figure S3). We then used the *lacZ* sequencing data from NGS and Sanger experiments in COSMIC format (Supplementary Figure S4) to identify which of the normalized signatures were most closely associated with the mutation spectrum of each agent. This initial analysis showed that mutational signatures in human cancers that have been associated with specific mutagenic exposures were enriched in the *lacZ* mutation profiles for the appropriate agent tested in this study (Figure 4). For example, the UVB^29, 31^ and sunlight^30^ mutation profiles had very strong correlations (Pearson’s coefficient = 0.93-0.98) with the SBS 7a signature, which is observed in human skin cancers. Similarly, the BaP mutation profile showed a strong correlation with several signatures including SBS 4 (Pearson’s coefficient = 0.76), which is observed in tobacco smoke-induced cancers. In total, there were six SBS signatures that had a Pearson’s coefficient greater than 0.8 with the mutation spectra generated from sequenced *lacZ* mutations (Figure 4).

**Figure 3.**
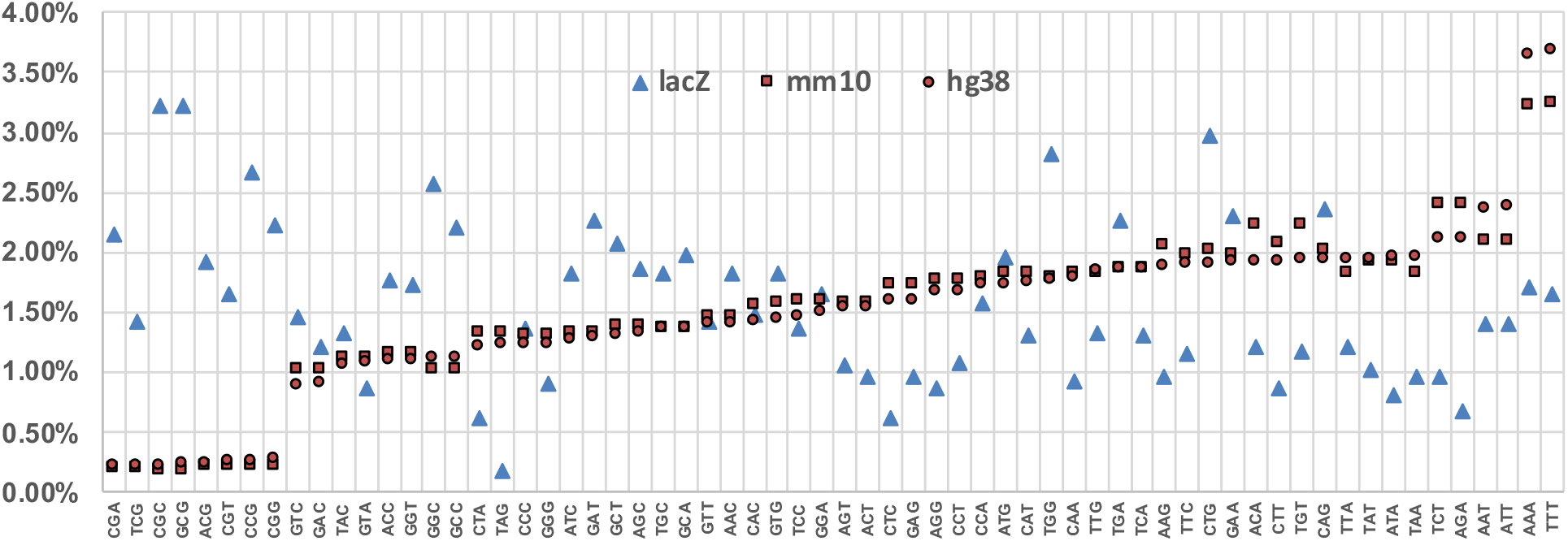
Trinucleotide context differences between the *lacZ* transgene, mouse genome, and human genome. Comparison of the frequencies of the 64 possible trinucleotides among the *lacZ* transgene (lacZ), mouse genome (mm10), and human genome (hg38) show that mouse and human genome frequencies are comparable with each other, while *lacZ* is more variable and biased towards some GC rich trinucleoties.

**Figure 4.**
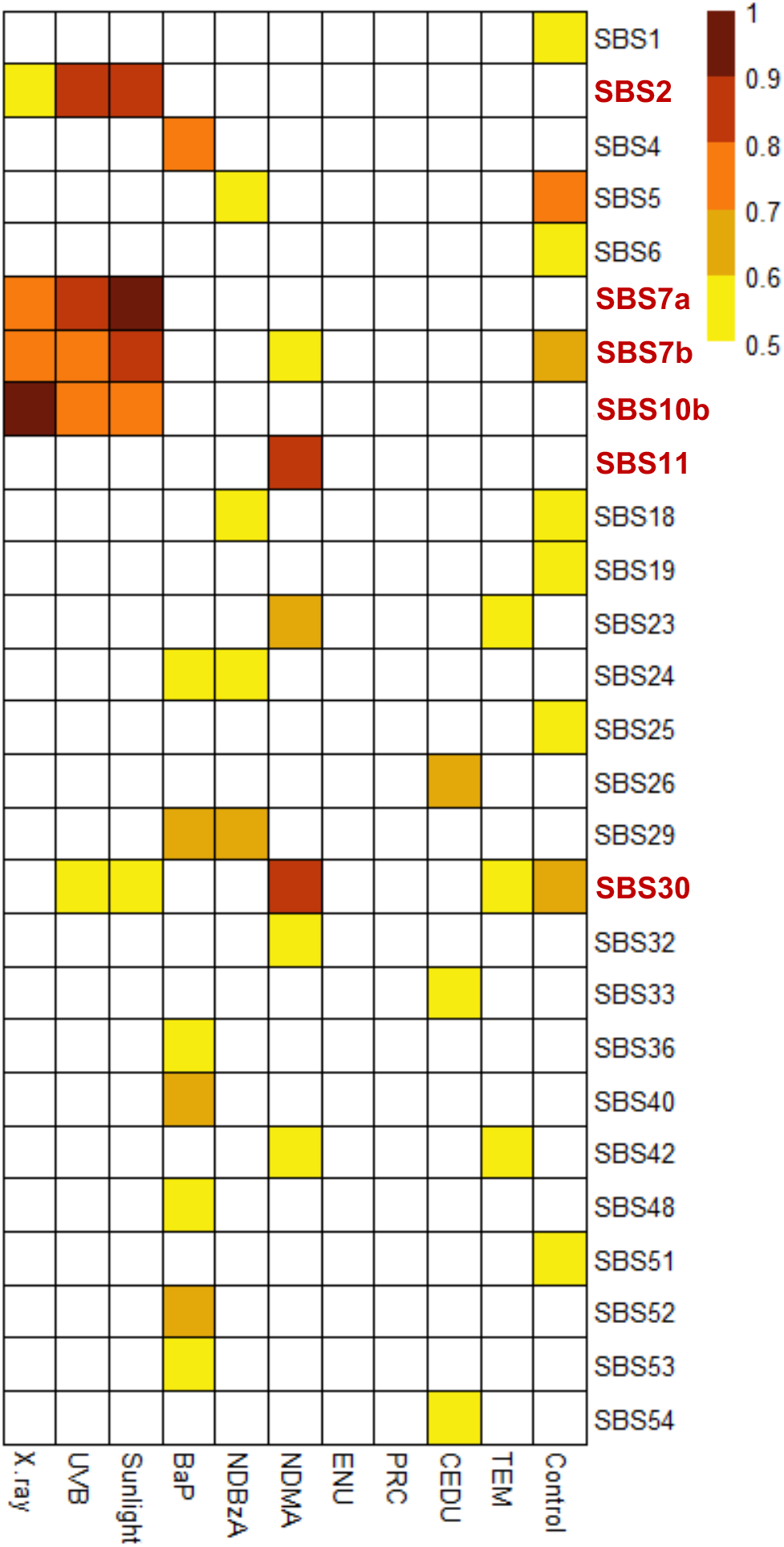
Heatmap of similarities between obtained mutational profiles of tested agents and COSMIC SBS signatures. All correlations that had a Pearson’s correlation above 0.5 are shown. The six SBS signatures that had a Pearson’s correlation greater than 0.7 are indicated in bold on the right of the heatmap.

Next, we used each of the agent-generated mutation data to simultaneously query the entire COSMIC SBS database to establish which of the signatures contributed to the observed spectra (Supplementary Figure S5). This process was conducted as described in steps 7-9 of Figure 1. Prior to this analysis, we used control mutation data to generate an *in vivo* background signature (Figure 5) to account for the fact that some mutations present in the exposure groups are also spontaneous in origin rather than specific to the mutagen tested. This is especially true for weak mutagens. As shown in Figure 5, the *in vivo* background signature is enriched primarily in C>T mutations, and to a lesser extent C>A mutations, and this was consistent among all tissues that contributed to the control signature (Supplementary Figure S6). Inclusion of the control signature improved the association between reconstructed signatures and mutation data by as much as 26% (Figure 6) and eliminated five of the weakly associated signatures (Supplementary Figure S7). Finally, application of stringent filtering criteria (see Methods) revealed the association of nine COSMIC SBS signatures with mutation data from the various exposure groups (Figure 6).

**Figure 5.**
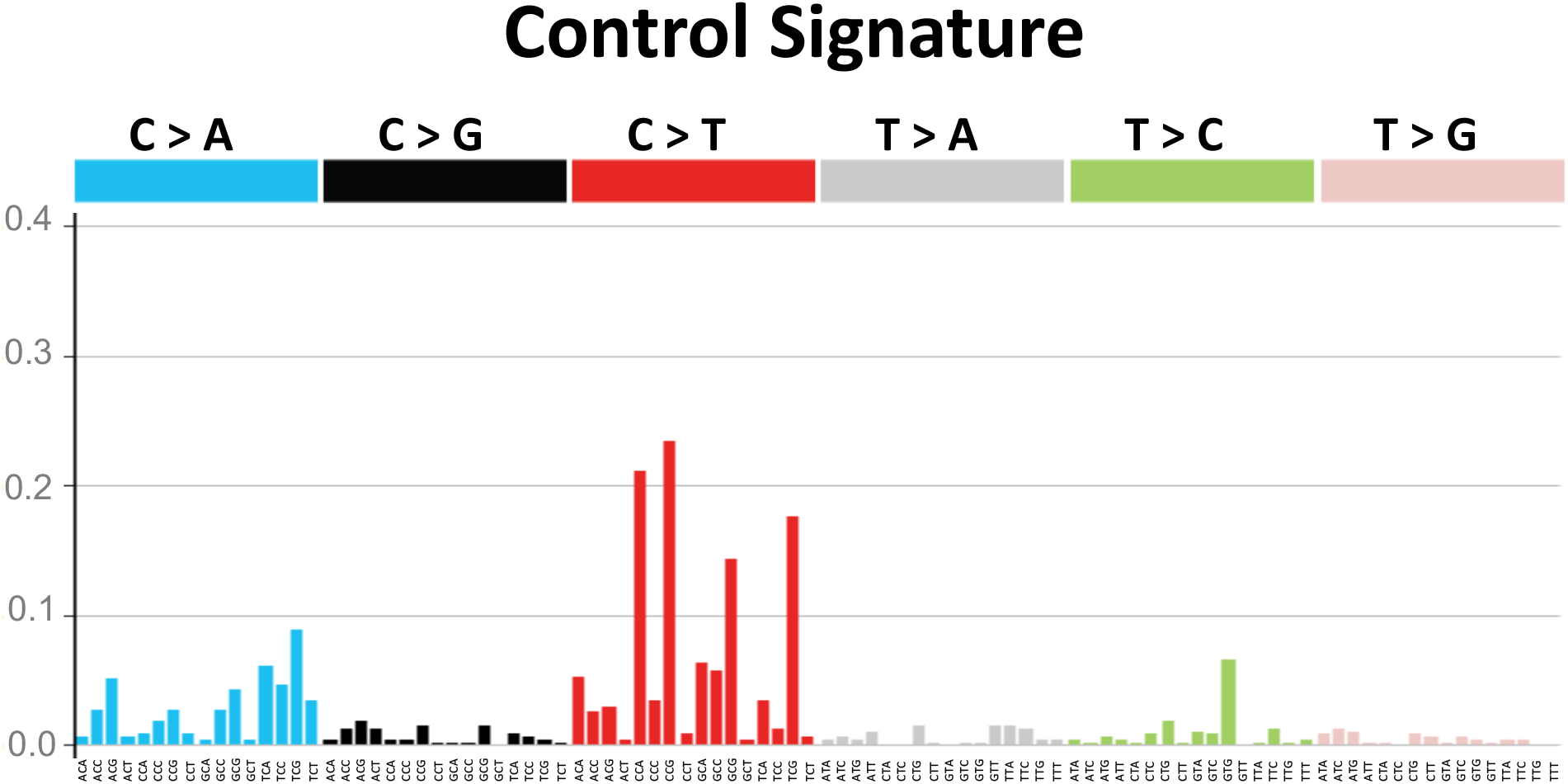
The *lacZ* control signature. The control signature is based on empirical mutation data from control animals in NGS and Sanger studies.

**Figure 6.**
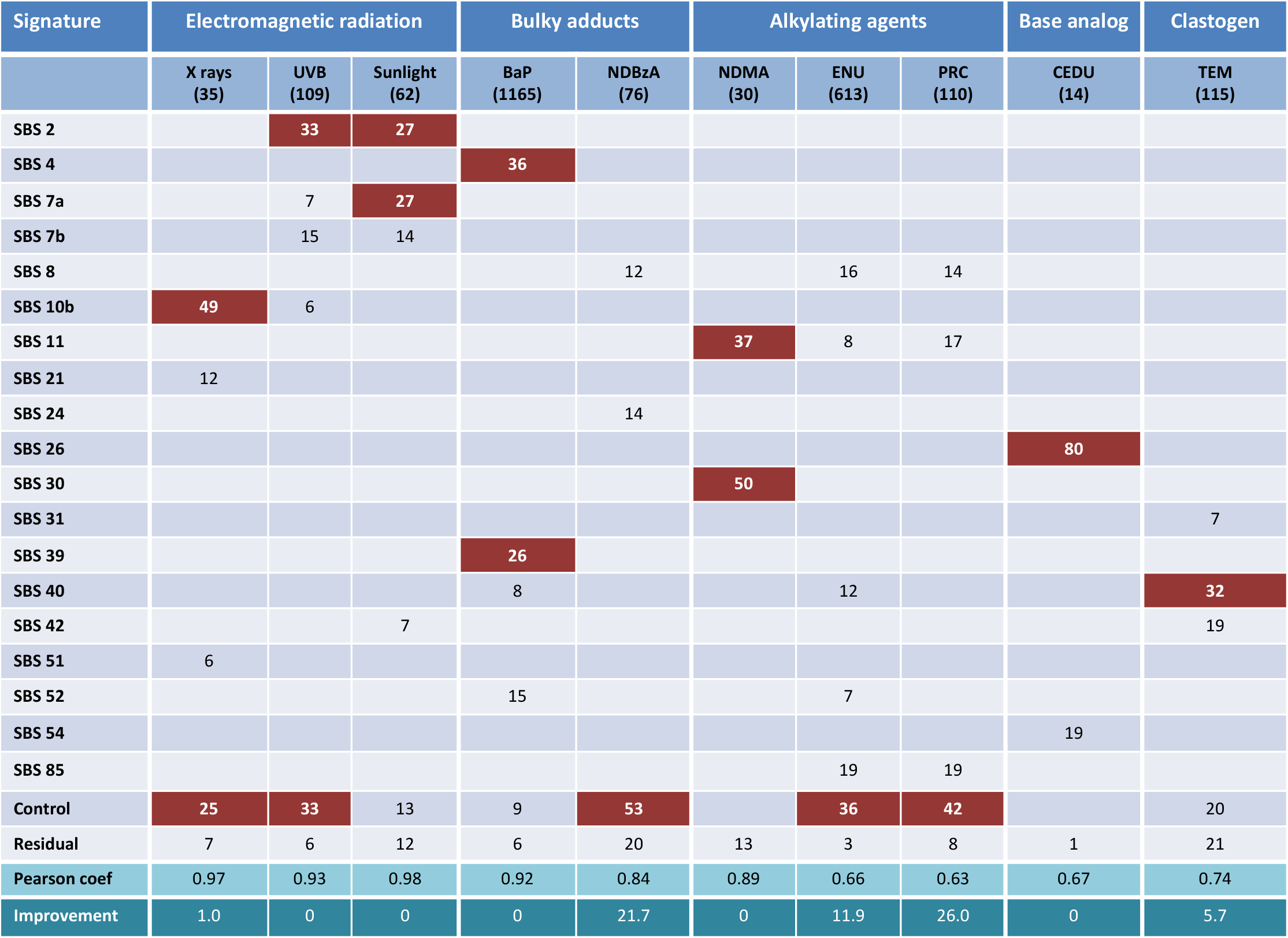
The association of mutational signatures across different exposure groups. The number below each agent indicates the number of unique mutants sequenced, while the number in each box represents the percent contribution of each signature to the mutation profile of each tested agent. Brown rectangles indicate that the signature was present in the characterized mutations following the mutagenic exposure and that this association was moderate to strong (Pearson’s coefficient > 0.5; other signatures i.e., with a coefficient <0.5). The last two rows report the Pearson’s coefficients between reconstructed signatures and observed mutation profiles and the percent increase due to inclusion of the control signature.

The signatures produced by the three electromagnetic radiations (i.e., UVB, sunlight, and X-rays) appear to be broadly similar when visually assessing individual SBS signature heatmaps (Figure 4). However, we found that different mutational processes contribute to each signature. Specifically, SBS 2 and the control signature each explained 33% of the UVB mutation profile; SBS 2 and SBS 7a each explained 27% of the sunlight data (Figure 6); and the mutation spectrum associated with X-rays, which induces large deletions rather than point mutations (56 indels ranging from 1-437 bp vs 35 SNVs^33^), was most associated with the SBS 10b signature (49%).

For the bulky adduct group, the mutation spectrum of BaP revealed mutational processes characteristic of SBS 4 (36%) and SBS 39 (27%) signatures. SBS 4 is most notably associated with tobacco-smoke induced cancer^16^, while SBS 39 is one of the new signatures that currently does not have a proposed etiology. No SBS signature was associated with the mutation profile of NDBzA and the control signature explained 53% of the mutation profile of this agent.

Analysis of the alkylating agent exposure group revealed that SBS 11 and SBS 30 signatures were associated with N-nitrosodimethylamine (NDMA) mutation data^34^ and explained 37% and 50% of the mutations, respectively. SBS 11 has previously been linked to exposures to the methylating agents temozolomide and N-methyl-N’-nitro-N-nitrosoguanidine^17,19^. SBS 30 is hypothesized to be associated with defects in base excision repair. No SBS signatures were associated with the mutation profiles of ENU or PRC while the control signature explained 36% and 42% of the mutation profiles of these two chemicals, respectively.

There were limited data available for nitrogenous base analogs. Data were only obtained from mice exposed to 5-(2-chloroethyl)-2-deoxyuridine (CEDU)^41^, a uridine analog. This included only 14 characterized mutants from bone marrow, 13 of which were T>C mutations. Consequently, 80% of the data were explained by the SBS 26 signature, which exhibits a bias for these types of substitutions.

TEM had a SNV mutation spectrum that was similar to controls (Figure 2). Nevertheless, we found that SBS 40 contributed to 32% of the TEM data, which was higher than the 20% that can be attributed to the control signature. There is currently no known etiology for the SBS 40 signature.

Overall, the reconstructed signatures had very strong Pearson’s coefficients (0.84 - 0.98) for six of the agents and strong coefficients (0.63 -0.74) for four agents with the respective *lacZ*-generated mutation profiles (Figure 6 and Supplementary Figure S4).

## DISCUSSION

We show that *in vivo* NGS-TGR data can be used to extract mutagenic mechanisms that may contribute to human cancers through application of COSMIC signature analysis. We also show that such analyses are improved through the inclusion of a background mutational signature (i.e., control signature) that reflects spontaneous mutations resulting from endogenous processes. Analysis of induced mutations in mouse tissues following exposures to 10 mutagenic agents (two sequenced by NGS, six sequenced by the Sanger method, and two by both) revealed high concordance between the expected mutagenic mode of action and the relevant COSMIC signature. The data suggest that our approach can be used to: (i) test if TGR mutation spectra support hypotheses that COSMIC signatures are attributed to particular mutagenic exposures, and (ii) generate hypotheses about the mutagenic mechanisms underlying human cancers through identifying enriched COSMIC signatures in TGR mutation spectra.

A large portion of mutations collected from weak mutagens are spontaneous rather than chemically induced. Thus, we developed a background signature derived from our empirical control data that can be integrated with COSMIC signatures to reduce the noise attributable to spontaneous mutation patterns. This *in vivo* control signature is a unique feature of our study, as there is currently no ‘background’ COSMIC signature and no *in vivo* control signature is reported in a recent study that generated chemical-specific signatures *in vivo* using a different approach^27^. Our results show that C>T transitions are the most common spontaneous mutations *in vivo* (Figure 5) and this was consistent among all tissues analyzed (Supplementary Figure 6). C>T transitions at CpG sites are known hotspots of mutation due to spontaneous deamination of cytosine^49^. Previous work using bisulfite sequencing has shown that CpG sites in *lacZ* are heavily methylated, and CpG flanked by a 5’ pyrimidine were most likely to have C>T base substitutions^46^. This is supported by our control data: the most prevalent spontaneous mutations were C>T at CCG, and, the third most prevalent were C>T mutations at T<u>C</u>G (Figure 5). Thus, our background control signature is consistent with expectations.

An *in vitro* background signature was recently reported^28^; however, the correlation between the two control signatures is modest (Pearson’s coefficient = 0.45) because, at variance with our results, the *in vitro* control signature is enriched for C>A mutations. Spontaneous deamination of cytosine is also the most likely reason for C>A transversions and seem to be the most common spontaneous mutation *in vitro*^53^. This suggests that the same type of event, i.e., cytosine deamination, can results in different outcomes, i.e., C>T versus C>A mutations, depending on the physiological context.

Application of the control signature (Figure 5) and stringent statistical analysis identified nine SBS signatures that were associated with the *lacZ* SNVs induced by the investigated exposures. Two major outcomes from this analysis are: 1) mutation profiles for some of the tested agents were highly enriched for COSMIC signatures from cancers where the agents are known etiological factors (e.g., UV for skin cancer and BaP for tobacco-related cancers); and, 2) a few *lacZ* mutation profiles were associated with a variety of signatures of unknown aetiologies. This raises the question of whether the mutagenic mechanisms of these prototype agents are determinants of the signatures.

We identified the SBS 2, SBS 7a, and SBS 10b signatures as important contributors to the mutagenic mechanisms of all three electromagnetic radiation agents investigated (i.e., X-ray, UVB, and sunlight). SBS 2 has been observed in ~14% of cancer samples and is present in 22 cancer types but is most often found in cervical and bladder cancers^14,17^. In this study, the signature was most strongly associated with UV skin exposure, representing 33-27% of mutations in exposed animals. Mechanistically, cytosine deamination is accelerated by UV exposure^54^; thus, it is possible that we observed SBS 2 in this study because of UV-dependent cytosine deamination. However, SBS 2 is not observed in skin cancers^17^. This suggests that mutations arising from UV-dependent cytosine deamination are not the primary drivers of the surveyed human skin cancers in the COSMIC database, and that other lesions (e.g., various types of photodimers) are the main contributors to the mutation catalogue of UV-induced skin cancers. Another possible explanation is that with a small sample size of mutations, the high degree of similarity in the SBS 2 and SBS 7a signatures confounds this analysis. By this logic, some portion of the mutational signature identified as SBS 2 in our study may be the result of the mutational processes associated with SBS 7a, which is found in multiple cancer types but is most pronounced in skin cancers^14,17^. Indeed, the SBS 7a signature contributes to 27% of the mutations observed after sunlight exposure.

Activation of error-prone polymerases has been attributed to SBS 10b^14^, a signature that is mostly found in colorectal and uterine cancers. In the present study, this signature was only associated with X-ray mutations (49%). X-ray mutations show a high proportion of C>T substitutions at the TCG motif (Supplementary Figure S4), which is characteristic of the *lacZ* normalized SBS 10b signature (Supplementary Figure S3). It is possible that there is an ionizing radiation component to this signature. However, given previous work in this area, it is more likely that the association between SBS 10b and X-ray SNVs is a result of error-prone replication occurring in response to DNA damage.

The analysis of mutational signatures for the electromagnetic radiation agents provide support for the ability of the expanded repertoire of COSMIC signatures to exploit subtle differences in the mutation profiles to extract different mutational mechanisms. Using the previous version of the COSMIC database, all three radiation types had a comparable contribution from signature 7 (21-33%; data not shown). However, there are now four SBS signatures (7a-7d) derived from the original signature 7 in the latest COSMIC database^15^, and of these, only the SBS 7a signature contributes significantly to the mutation profile of sunlight.

Tobacco smoking is strongly associated with SBS 4, and this signature is commonly found in the lung tumors of smokers. BaP is a major mutagenic component in tobacco smoke^21^ and as expected, SBS 4 contributed the highest percentage (36%) to the mutation profile of BaP. Interestingly, SBS4 was the only signature that contributed to the mutation profile of BaP and accounted for 60% of the observed profile when using the previous version of the COSMIC database (data not shown). However, using version 3 of the COSMIC database^15^, the contribution of SBS 4 declined while we identified a second signature that contributed to the BaP mutation profile. Specifically, we detected a significant contribution (27%) of the SBS 39 signature, which is one of the new signatures and currently has no known etiology. These results suggest an overlap in the aetiology of these two signatures and that SBS 39 may be associated with exposure to chemicals that induce bulky adducts.

The BaP mutation profile that we derived using our approach is consistent with previous work *in vivo*^27^ and *in vitro*^28^ that demonstrated the presence of SBS 4 after exposure to BaP. Indeed, the BaP mutation profile is consistent among the three studies (Pearson’s correlation of 0.80 and 0.71 with the *in vivo* and *in vitro* profile, respectively). Remarkably, signatures SBS 24, which has been associated with aflatoxin adducts, and SBS 29, which has been associated with tobacco chewing, are strikingly similar to SBS 4 (Supplementary Figure S3). However, only SBS 4 strongly correlates with the BaP mutation data. This demonstrates the robustness of the mutational signatures and the ability of TGR-NGS to correctly discriminate between similar signatures that have different aetiologies. It also emphasizes the importance of the flanking nucleotides to increasing the specificity of the signatures; this work demonstrates that 96-bp signatures provide superior mechanistic information to standard mutation spectrum analysis.

NDMA was the only alkylating agent among those investigated that was associated with an established COSMIC signature. About 50% of the NDMA mutation profile was explained by the SBS 30 signature that has been associated with a deficiency in base excision repair. NDMA is known to induce mostly 0^6^- and N^7^-methyl guanine adducts^34^, thus, a role of base excision repair in the response to this chemical is expected. NDMA exposure was also enriched for SBS 11 (37%), inducing primarily C>T mutations at <u>C</u>pC motifs (Supplementary Figure S4). SBS 11 has been detected in melanomas and glioblastomas, and the mutation pattern of this signature has been attributed to alkylating agent exposures, such as temozolomide and N-methyl-N’-nitro-N-nitrosoguanidine^17,19^. These alkylating agents induce C>T mutations, mostly at <u>C</u>pC motifs, and mutations at this motif are the four most common in the SBS 11 signature. The TGR mutation data from our study are consistent with this expected mutation spectrum.

The SBS 11 signature was not enriched within the mutation spectrum of the two other alkylating agents (i.e., ENU or PRC) in our mutation database. This is expected because these compounds induce a very different mutation spectrum, causing primarily T>A mutations. These differences demonstrate that SBS 11 is specific to a particular mechanism of alkylation (i.e., target sites for the alkylation events) and that there is currently no COSMIC signature for alkylating agents that target thymine. Further TGR-NGS analyses of alkylating agents may refine our understanding regarding which specific alkylating agents or defective alkyltransferases underlie the mechanisms associated with SBS 11.

The mutation profile obtained with ENU, demonstrating a slight preponderance of T>A mutations over T>C mutations, is consistent (Pearson’s coefficient = 0.71) with that obtained in the bone marrow of *gpt* delta mice^27^, although the correlation is reduced when expanding the six possible base pair alterations to the 96 possible mutation types (Pearson’s coefficient = 0.49). This is mostly due to a deficiency of T>C mutations at CTN motifs with respect to *gpt* delta mice. Nevertheless, the similarity with the ENU mutation profile from *gpt* delta mice is greater than that obtained *in vitro* with an induced pluripotent stem cell (iPSC) line (Pearson’s coefficient = 0.30) where the ENU signature is dominated by T>C mutations^28^. These authors speculate that the preponderance of T>C mutations after *in vitro* exposure to ENU is driven by the intrinsic characteristics of DNA repair processes in iPSCs.

The mutation profile of CEDU, a nitrogenous base analog, closely matched the SBS 26 signature, which contributed to 80% of mutations in exposed animals. This is the highest contribution of a COSMIC signature to any of the mutation profiles generated in this study. SBS 26 is one of the seven SBS signatures associated with defective mismatch repair, which is one of the major repair pathways that deals with base analogs^55^. Due to the limited number of mutations recovered in the CEDU study, the association between SBS 26 and CEDU should be further tested. Also, considering that CEDU is similar in structure to existing halogenated uracil analogs that serve as therapeutics (e.g., fluorouracil), attention should be given to these compounds as possible contributors to the SBS 26 signature and associated cancers.

Among the agents tested in this study, TEM is the only one that is more effective at inducing chromosomal structural aberrations than mutations. TEM is a trifunctional alkylating agent that induced a strong micronucleus response while eliciting a weak mutagenic response in the hematopoietic system^48^. Our analysis identified SBS 40 signature as a strong contributor (32%) to the mutation profile of TEM. SBS 40 is one of those signatures that is not dominated by any specific type of base pair alteration and does not have a proposed aetiology. Further studies are needed to confirm whether SBS 40 signature is an indicator of a clastogenic mode of action.

In summary, we demonstrated that *lacZ* transgene sequence data can be used, in conjunction with established mutation signatures derived from COSMIC cancer data sets, to test the hypothesis that a given class of mutagenic agents is linked with specific human cancers. Moreover, COSMIC signature mining based on TGR mutation data sets can be used to generate new hypotheses regarding the mutagenic mechanisms associated with human cancers. This study presents a potential avenue through which mutation signature analysis can be applied to *in vivo* experimental models, and the analyses employed to improve understanding of mode of action. The analyses can also generate hypotheses regarding the mutational mechanisms of uncharacterized chemicals. The *in vivo* TGR-NGS approach has comparable sensitivity to whole genome approaches used for investigating the mutational landscape of environmental agents^18, 19, 26, 28, 56^. However, by avoiding the orders-of-magnitude higher cost of whole-genome sequencing, the *in vivo* TGR-NGS approach offers much higher-throughput for the testing of chemical mutagens. Overall, these results highlight that some mutational signatures may have large environmental components and contribute to the growing body of evidence that analyses of mutations spectra shortly after exposure has bearing on the carcinogenic mechanism and the mutational profile observed in fully developed cancers.

## MATERIALS AND METHODS

### Animal Treatment

Male MutaMouse animals (6-15 weeks old; 6-8 per group) were exposed daily to either 100 mg/kg BaP, 5 mg/kg ENU, 25 mg/kg PRC or 2 mg/kg TEM by oral gavage for 28 days as per the Organisation for Economic Co-operation and Development (OECD) test guideline 488^57^. All doses were selected based on pilot studies conducted to identify the maximum tolerated dose as per TG 488 guidance. The BaP^8^, PRC^47^ and TEM^48^ data are the same presented their respective reference. Matched controls received the solvent (olive oil or water) by oral gavage during the same period. Three days after the last daily exposure, mice were anaesthetized with isofluorane and euthanized via cervical dislocation. Bone marrow cells were isolated by flushing femurs with 1X phosphate-buffered saline. After brief centrifugation, the supernatant was discarded, and the pellet was flash-frozen in liquid nitrogen prior to storage at −80 °C. All animal procedures were carried out under conditions approved by the Health Canada Ottawa Animal Care Committee.

### *lacZ* Mutant Quantification, Collection, and Sequencing

The experimental protocol for enumerating *lacZ* mutants followed OECD guideline 488^57^. Briefly, bone marrow was thawed and digested overnight with gentle shaking at 37 °C in 5 mL of lysis buffer (10 mM Tris–HCl, pH 7.6, 10 mM ethylenediaminetetraacetic acid (EDTA), 100 mM NaCl, 1 % sodium dodecyl sulfate (w/v), 1 mg/mL Proteinase K). High molecular weight genomic DNA was isolated using phenol/chloroform extraction as described previously^42^,^58^. The isolated DNA was dissolved in 100 μL of TE buffer (10 mM Tris pH 7.6, 1 mM EDTA) and stored at 4 °C for several days before use. The phenyl-β-D-galactopyranoside (P-gal) positive selection assay^59^ was used to identify *lacZ* mutants present in the DNA. Briefly, the λgt10lacZ construct present in the genomic DNA was isolated and packaged into phage particles using the Transpack™ lambda packaging system (Agilent, Mississauga, Ontario, Canada). The phages were then mixed with *E. coli* (*lacZ^−^*, *galE^−^*, *recA^−^*, *pAA119^−^* with *galT* and *galK*)^58^ in order to transfect the cells with the *lacZ* construct. *E. coli* were then plated on a selective media containing 0.3% P-gal (w/v) and incubated overnight at 37 °C. Only *E. coli* receiving a mutant copy of *lacZ* where the gene function is disrupted can form plaques on the P-gal medium, because P-gal is toxic to *galE^−^* strains with a functional *lacZ* gene product^1^. Packaged phage particles were concurrently plated on plates without P-gal (titre plates) to quantify the total plaque-forming units to be used as the denominator in the mutant frequency calculation.

After enumeration, plaques from each individual sample were collected and pooled together in microtubes containing autoclaved milliQ water (0.3 plaques/μL; mutants from 1 sample per tube). Mutant amplification and sequencing were done as described previously^8^. Briefly, the mutant pools were boiled for 5 minutes and transferred to a PCR mastermix containing a final concentration of 1X Q5 reaction buffer, 200 μM dNTPs, 0.5 μM Forward primer (GGCTTTACACTTTATGCTTC), 0.5 μM Reverse Primer (ACATAATGGATTTCCTTACG), and 1U Q5 enzyme (New England BioLabs Ltd., Whitby, Ontario, Canada); the final volume of each PCR was 50 μL. To control for errors introduced during PCR, each mutant pool was amplified twice as two separate technical replicates. The following thermocycle program was used for amplification: 95 °C for 3 min; 30 cycles of 95 °C for 45 s, 50 °C for 1 min, 72 °C for 4 min; final extension at 72 °C for 7 min. PCR products were purified using the QIAquick PCR purification kit (Qiagen, Montreal, Quebec, Canada).

NGS libraries were built using the NEBNext® Fast DNA Library Prep Set for Ion Torrent™. Each technical replicate had a unique barcoded adaptor ligated to the *lacZ* DNA fragments allowing for many samples to be sequenced simultaneously (up to 96 libraries per NGS run). Sequencing was performed using the Ion Chef™ workflow and Ion Proton^™^ system with P1 chips. NGS reads were aligned to the *lacZ* gene using bowtie 2 (version 2.1.0) and read depths for every possible mutation were quantified using samtools (version 0.1.19). Mutations were called if, after background correction (determined by sequencing non-mutants), both technical replicates had mutation read depths above threshold values (equal to at least 1/number of plaques in pool)^8^. To further filter the data in this study, if the mutation read depths between two technical replicates varied by ≥50% then that mutation was removed from analysis. Clonally expanded mutants were only counted as one mutation.

### Published Sanger Sequencing Data

Published data came from studies where *lacZ* transgene mutants were sequenced and the position and type of each mutation was reported (summarized in Supplementary Table S1). Mutants were characterized from MutaMouse or *LacZ* Plasmid mice^60^. Some studies reported the position of the mutation in the plasmid construct, while others reported the position in the coding sequence. For consistency, the positional information was adjusted to reflect the position of the mutation in the coding sequence of the *lacZ* gene. Furthermore, the reference sequence of *lacZ* used for NGS has four variations^38^ relative to the *E. coli lacZ* coding sequence (Genbank: V00296.1)^61^, including a 15 bp insertion into codon 8. Thus, mutation positions were also adjusted to reflect this where applicable (e.g., if *LacZ* Plasmid mice were used instead of MutaMouse). No mutations were detected at or next to the variant positions in the *LacZ* Plasmid motif. In contrast to NGS work, different tissues were used for these analyses (i.e., bone marrow, brain, colon, germ cells, kidney, liver, skin, spleen, and stomach). Tissue sources are noted in the results with the accompanying data.

### Signature Analyses

The workflow used to do signature analyses are available as an RShiny web-application (https://github.com/MarcBeal/HC-MSD/tree/master/lacZ_Mutations_COSMIC_Signatures). Mutations for control and exposed samples (see metadata in Supplemental Material) were imported into the R console^62^ as VRanges using the package “VariantAnnotation”^63^ with the *lacZ* coding sequence as the reference FASTA file. To determine which of the COSMIC mutation signatures best explained the observed *lacZ* mutant spectrum, the COSMIC mutation signature weights, which are derived from human mutation data, were first normalized to *lacZ* trinucleotide frequencies. This was done using the ratio of trinucleotide frequencies in *lacZ* to the trinucleotide frequencies in the human genome (Figure 3; the normalized signatures are shown in Supplementary Figure S3 and the raw numbers in Supplementary Material). Analysis was done this way (as opposed to converting *lacZ* mutation data themselves to human trinucleotide frequencies) because the COSMIC signature are based on a much larger database, and therefore, represent a more robust signal with less variance. Following normalization, each of the 96 trinucleotide substitutions within each signature were represented as the relative frequency (i.e., all values in a signature sum to 1) by dividing each normalized value by the sum of all values for that signature. The trinucleotide mutation context (i.e., the nucleotide immediately upstream and downstream of the mutation) was obtained with the “mutationContext” function and converted to a motif matrix using the “motifMatrix” function (both in the “SomaticSignatures” package^64^). The motif matrix was then transposed to obtain the required format, and finally decomposed into the constituent *lacZ*-normalized signatures using the “whichSignatures” function from “deconstructSigs”^65^. The contribution of each identified signature to the mutation data was reported as a fraction. If the sum of each signature did not account for 100% of the mutation data, then the remainder was reported as the “residual”.

In order to account for spontaneous mutations often present alongside induced mutations, which is especially true for weak mutagens, we generated a signature for the spontaneous mutation background using the mutations observed in control animals. This included all control mutations characterized by NGS and Sanger sequencing. However, spontaneous SNVs characterized by Sanger sequencing were heavily biased towards positions 1072, 1090, 1187, 1627, and 2374. Therefore, Sanger sequencing data at these 5 positions were not used for deriving the control mutation signature. Signatures were plotted using ggplot2^66^.

“Signature reconstruction” was then used to determine how well the combination of normalized signatures, identified using the “whichSignatures” function, explain the mutation data from the respective exposure groups. For example, if signatures 3 and 4 contributed 40% and 60% to the mutation profile of a compound, respectively, then the motif matrices for signatures 3 and 4 were multiplied by 0.4 and 0.6, respectively, and summed together. The reconstructed signature was then compared against the motif matrices of the compound using Pearson correlation.

Lastly, the contribution of individual signatures was further validated using Pearson correlation. Specifically, each signature was compared against the respective 96-base context mutation spectra from which the signature was identified. In the final results, COSMIC signatures were only reported if the contribution was greater than the largest residual, and the Pearson coefficient with the reconstructed signature was greater than 0.5.

### Statistics

Statistical analyses were done using the R programming language^62^. Mutant frequencies were compared between exposure groups and controls using generalized estimating equations assuming a Poisson distribution for the error, as done previously^8^, using the geepack library^67^ with outliers (1 in control, 1 in TEM) removed. Bonferonni correction for multiple comparisons was used to adjust the threshold of significance. Mutation spectra of the chemical exposure groups were compared against controls using mutation proportions. The standard error for the mutation spectra was determined using error propagation. Significant differences in mutation spectra between chemically induced mutants and spontaneous control mutants were determined using Fisher’s exact tests with Bonferroni correction for multiple comparisons (i.e., across different chemical groups). To compare whole mutation spectra between control and exposed groups, Fisher’s exact tests were performed with Monte Carlo simulation with 10,000 replicates. Fisher’s exact tests were also performed on 2 × 2 sub-tables for each mutation type.

## ACKNOWLEDGEMENTS

We would like to thank Angela Dykes, Lynda Soper, and John Gingerich for their technical contributions to this research. We are grateful for advice provided by Dr. Ludmil Alexandrov and Andrew Williams.

## AUTHOR CONTRIBUTIONS

MAB, CM, MJM and JOB conducted the MutaMouse animal studies and collected samples. MAB and MJM sequenced plaques. MAB, MJM and DL conducted the COSMIC analyses. CY and FM secured funding for the study and were responsible for study conception and design. All authors contributed to data analysis, interpretation, paper writing and approved the final version of the manuscript.

## References

1. Lambert IB, Singer TM, Boucher SE, Douglas GR. Detailed review of transgenic rodent mutation assays. Mutat Res 590, 1–280 (2005).

2. OECD. Detailed Review Paper on Transgenic Rodent Mutations Assay, Paris (2009).

3. Meier MJ, Beal MA, Schoenrock A, Yauk CL, Marchetti F. Whole genome sequencing of the Mutamouse model reveals strain- and colony-level variation, and genomic features of the transgene integration site. Sci Rep 9, 13775 (2019).

4. Shwed PS, Crosthwait J, Douglas GR, Seligy VL. Characterisation of MutaMouse lambdagt10-lacZ transgene: evidence for in vivo rearrangements. Mutagenesis 25, 609–616 (2010).

5. Gingerich JD, Soper L, Lemieux CL, Marchetti F, Douglas GR. Transgenic Rodent Gene Mutation Assay in Somatic Tissues. Springer Science+Business Media (2014).

6. O’Brien JM, et al. Transgenic rodent assay for quantifying male germ cell mutant frequency. J Vis Exp, e51576 (2014).

7. Besaratinia A, Li H, Yoon JI, Zheng A, Gao H, Tommasi S. A high-throughput next-generation sequencing-based method for detecting the mutational fingerprint of carcinogens. Nucleic Acids Res 40, e116 (2012).

8. Beal MA, Gagne R, Williams A, Marchetti F, Yauk CL. Characterizing benzo[a]pyrene-induced lacZ mutation spectrum in transgenic mice using next-generation sequencing. BMC Genomics 16, 812 (2015).

9. Meier MJ, O’Brien JM, Beal MA, Allan B, Yauk CL, Marchetti F. In utero exposure to benzo[a]pyrene increases mutation burden in the soma and sperm of adult mice. Environ Health Perspect 125, 82–88 (2017).

10. Alexandrov LB, Nik-Zainal S, Wedge DC, Campbell PJ, Stratton MR. Deciphering signatures of mutational processes operative in human cancer. Cell Rep 3, 246–259 (2013).

11. Nik-Zainal S, et al. Mutational processes molding the genomes of 21 breast cancers. Cell 149, 979–993 (2012).

12. Nik-Zainal S, et al. The life history of 21 breast cancers. Cell 149, 994–1007 (2012).

13. Forbes SA, et al. COSMIC: somatic cancer genetics at high-resolution. Nucleic Acids Res 45, D777–D783 (2017).

14. Helleday T, Eshtad S, Nik-Zainal S. Mechanisms underlying mutational signatures in human cancers. Nat Rev Genet 15, 585–598 (2014).

15. Alexandrov LB, et al. The Repertoire of Mutational Signatures in Human Cancer. BioRxiv, (2018).

16. Alexandrov LB, et al. Mutational signatures associated with tobacco smoking in human cancer. Science 354, 618–622 (2016).

17. Alexandrov LB, et al. Signatures of mutational processes in human cancer. Nature 500, 415–421 (2013).

18. Nik-Zainal S, et al. The genome as a record of environmental exposure. Mutagenesis 30, 763–770 (2015).

19. Olivier M, et al. Modelling mutational landscapes of human cancers in vitro. Sci Rep 4, 4482 (2014).

20. Phillips DH. Mutational spectra and mutational signatures: Insights into cancer aetiology and mechanisms of DNA damage and repair. DNA Repair (Amst) 71, 6–11 (2018).

21. Pfeifer GP, Denissenko MF, Olivier M, Tretyakova N, Hecht SS, Hainaut P. Tobacco smoke carcinogens, DNA damage and p53 mutations in smoking-associated cancers. Oncogene 21, 7435–7451 (2002).

22. Alexandrov LB, et al. Clock-like mutational processes in human somatic cells. Nat Genet 47, 1402–1407 (2015).

23. Hollstein M, Alexandrov LB, Wild CP, Ardin M, Zavadil J. Base changes in tumour DNA have the power to reveal the causes and evolution of cancer. Oncogene 36, 158–167 (2017).

24. Zhivagui M, Korenjak M, Zavadil J. Modelling mutation spectra of human carcinogens using experimental systems. Basic Clin Pharmacol Toxicol 121 Suppl 3, 16–22 (2017).

25. Chawanthayatham S, et al. Mutational spectra of aflatoxin B1 in vivo establish biomarkers of exposure for human hepatocellular carcinoma. Proc Natl Acad Sci U S A 114, E3101–E3109 (2017).

26. Huang MN, et al. Genome-scale mutational signatures of aflatoxin in cells, mice, and human tumors. Genome Res 27, 1475–1486 (2017).

27. Matsumura S, Sato H, Otsubo Y, Tasaki J, Ikeda N, Morita O. Genome-wide somatic mutation analysis via Hawk-Seq reveals mutation profiles associated with chemical mutagens. Arch Toxicol 93, 2689–2701 (2019).

28. Kucab JE, et al. A Compendium of Mutational Signatures of Environmental Agents. Cell 177, 821–836 e816 (2019).

29. Ikehata H, Masuda T, Sakata H, Ono T. Analysis of mutation spectra in UVB-exposed mouse skin epidermis and dermis: frequent occurrence of C- - >T transition at methylated CpG-associated dipyrimidine sites. Environ Mol Mutagen 41, 280–292 (2003).

30. Ikehata H, Nakamura S, Asamura T, Ono T. Mutation spectrum in sunlight-exposed mouse skin epidermis: small but appreciable contribution of oxidative stress-mediated mutagenesis. Mutat Res 556, 11–24 (2004).

31. Frijhoff AF, et al. UVB-induced mutagenesis in hairless lambda lacZ-transgenic mice. Environ Mol Mutagen 29, 136–142 (1997).

32. Ono T, Ikehata H, Vishnu Priya P, Uehara Y. Molecular nature of mutations induced by irradiation with repeated low doses of X-rays in spleen, liver, brain and testis of lacZ-transgenic mice. Int J Radiat Biol 79, 635–641 (2003).

33. Ono T, et al. Molecular nature of mutations induced by a high dose of x-rays in spleen, liver, and brain of the lacZ-transgenic mouse. Environ Mol Mutagen 34, 97–105 (1999).

34. Souliotis VL, van Delft JH, Steenwinkel MJ, Baan RA, Kyrtopoulos SA. DNA adducts, mutant frequencies and mutation spectra in lambda lacZ transgenic mice treated with N-nitrosodimethylamine. Carcinogenesis 19, 731–739 (1998).

35. Suzuki T, et al. A comparison of the genotoxicity of ethylnitrosourea and ethyl methanesulfonate in lacZ transgenic mice (Muta Mouse). Mutat Res 395, 75–82 (1997).

36. Mientjes EJ, et al. DNA adducts, mutant frequencies, and mutation spectra in various organs of lambda lacZ mice exposed to ethylating agents. Environ Mol Mutagen 31, 18–31 (1998).

37. Jiao J, Douglas GR, Gingerich JD, Soper LM. Analysis of tissue-specific lacZ mutations induced by N-nitrosodibenzylamine in transgenic mice. Carcinogenesis 18, 2239–2245 (1997).

38. Hakura A, Tsutsui Y, Sonoda J, Tsukidate K, Mikami T, Sagami F. Comparison of the mutational spectra of the lacZ transgene in four organs of the MutaMouse treated with benzo[a]pyrene: target organ specificity. Mutat Res 447, 239–247 (2000).

39. Douglas GR, Jiao J, Gingerich JD, Gossen JA, Soper LM. Temporal and molecular characteristics of mutations induced by ethylnitrosourea in germ cells isolated from seminiferous tubules and in spermatozoa of lacZ transgenic mice. Proc Natl Acad Sci U S A 92, 7485–7489 (1995).

40. Douglas GR, Jiao J, Gingerich JD, Soper LM, Gossen JA. Temporal and molecular characteristics of lacZ mutations in somatic tissues of transgenic mice. Environ Mol Mutagen 28, 317–324 (1996).

41. Staedtler F, Suter W, Martus HJ. Induction of A:T to G:C transition mutations by 5-(2-chloroethyl)-2’-deoxyuridine (CEDU), an antiviral pyrimidine nucleoside analogue, in the bone marrow of Muta Mouse. Mutat Res 568, 211–220 (2004).

42. Douglas GR, Gingerich JD, Gossen JA, Bartlett SA. Sequence spectra of spontaneous lacZ gene mutations in transgenic mouse somatic and germline tissues. Mutagenesis 9, 451–458 (1994).

43. Dolle ME, Martus HJ, Novak M, van Orsouw NJ, Vijg J. Characterization of color mutants in lacZ plasmid-based transgenic mice, as detected by positive selection. Mutagenesis 14, 287–293 (1999).

44. Dolle ME, Snyder WK, Dunson DB, Vijg J. Mutational fingerprints of aging. Nucleic Acids Res 30, 545–549 (2002).

45. Dolle ME, et al. Increased genomic instability is not a prerequisite for shortened lifespan in DNA repair deficient mice. Mutat Res 596, 22–35 (2006).

46. Ikehata H, Takatsu M, Saito Y, Ono T. Distribution of spontaneous CpG-associated G:C - - > A:T mutations in the lacZ gene of Muta mice: effects of CpG methylation, the sequence context of CpG sites, and severity of mutations on the activity of the lacZ gene product. Environ Mol Mutagen 36, 301–311 (2000).

47. Maurice C, Dertinger SD, Yauk CL, Marchetti F. Integrated In Vivo Genotoxicity Assessment of Procarbazine Hydrochloride Demonstrates Induction of Pig-a and LacZ Mutations, and Micronuclei, in MutaMouse Hematopoietic Cells. Environ Mol Mutagen 60, 505–512 (2019).

48. Maurice C, O’Brien JM, Yauk CL, Marchetti F. Integration of sperm DNA damage assessment into OECD test guidelines for genotoxicity testing using the MutaMouse model. Toxicol Appl Pharmacol 357, 10–18 (2018).

49. Duret L. Mutation patterns in the human genome: more variable than expected. PLoS Biol 7, e1000028 (2009).

50. Shelby MD, Tindall KR. Mammalian germ cell mutagenicity of ENU, IPMS and MMS, chemicals selected for a transgenic mouse collaborative study. Mutat Res 388, 99–109 (1997).

51. Beranek DT. Distribution of methyl and ethyl adducts following alkylation with monofunctional alkylating agents. Mutat Res 231, 11–30 (1990).

52. Revollo J, et al. Spectrum of Pig-a mutations in T lymphocytes of rats treated with procarbazine. Mutagenesis 32, 571–579 (2017).

53. de Jong PJ, Grosovsky AJ, Glickman BW. Spectrum of spontaneous mutation at the APRT locus of Chinese hamster ovary cells: an analysis at the DNA sequence level. Proc Natl Acad Sci U S A 85, 3499–3503 (1988).

54. Peng W, Shaw BR. Accelerated deamination of cytosine residues in UV-induced cyclobutane pyrimidine dimers leads to CC- - >TT transitions. Biochemistry 35, 10172–10181 (1996).

55. Kunkel TA. DNA-mismatch repair. The intricacies of eukaryotic spell-checking. Curr Biol 5, 1091–1094 (1995).

56. Meier B, et al. C. elegans whole-genome sequencing reveals mutational signatures related to carcinogens and DNA repair deficiency. Genome Res 24, 1624–1636 (2014).

57. OECD. Test 488: Transgenic Rodent Somatic and Germ Cells Gene Mutation Assays. OECD Publishing (2013).

58. Gossen JA, Molijn AC, Douglas GR, Vijg J. Application of galactose-sensitive E. coli strains as selective hosts for LacZ-plasmids. Nucleic Acids Res 20, 3254 (1992).

59. Vijg J, Douglas GR. Bacteriophage lambda and plasmid lacZ transgenic mice for studying mutations in vivo. In: Technologies for detection of DNA damage and mutations (ed^(eds Pfeifer GP). Plenum Press (1996).

60. Vijg J, Dolle ME, Martus HJ, Boerrigter ME. Transgenic mouse models for studying mutations in vivo: applications in aging research. Mech Ageing Dev 99, 257–271 (1997).

61. Kalnins A, Otto K, Ruther U, Muller-Hill B. Sequence of the lacZ gene of Escherichia coli. EMBO J 2, 593–597 (1983).

62. R Core Team. R: a language and environment for statistical computing (2016).

63. Obenchain V, Lawrence M, Carey V, Gogarten S, Shannon P, Morgan M. VariantAnnotation: a Bioconductor package for exploration and annotation of genetic variants. Bioinformatics 30, 2076–2078 (2014).

64. Gehring JS, Fischer B, Lawrence M, Huber W. SomaticSignatures: inferring mutational signatures from single-nucleotide variants. Bioinformatics 31, 3673–3675 (2015).

65. Rosenthal R, McGranahan N, Herrero J, Taylor BS, Swanton C. DeconstructSigs: delineating mutational processes in single tumors distinguishes DNA repair deficiencies and patterns of carcinoma evolution. Genome Biol 17, 31 (2016).

66. Wickham H. ggplot2: elegant graphics for data analysis. Springer-Verlag (2016).

67. Halekoh U, Højsgaard S, Yan J. The R package geepack for generalized estimating equations. Journal of Statistical Software 15, 1–11 (2006).

