## Supplementary tables and figures for "Sequencing Chemically Induced Mutations in the Mutamouse Lacz Reporter Gene Identifies Human Cancer Mutational Signatures"

[1]Environmental Health Science and Research Bureau, Healthy Environments and Consumer Safety Branch, Health Canada, Ottawa, Ontario, K1A 0K9, Canada.

[2]Science and Technology Branch, Environment and Climate Change Canada, Ottawa, Ontario, K1A 0H3, Canada.

[3]National Wildlife Research Centre, Environment and Climate Change Canada, Ottawa, Ontario, K1A 0H3, Canada.

[4]Present address: Existing Substances Risk Assessment Bureau, Health Canada, Ottawa, Ontario, Canada

Running title: Induced lacZ mutations and human cancer signatures

### SUPPLEMENTARY TABLES AND FIGURES

**Supplementary Table S1.** Summary of published data from studies that used Sanger sequencing to characterize *lacZ* mutants.

| Chemical | Tissues | Total Mutations | SNVs | References |
|---|---|---|---|---|
| BaP | Colon, Spleen, Stomach | 78 | 60 | (27) |
| CEDU | Bone Marrow | 14 | 14 | (30) |
| ENU | Bone Marrow, Germ Cells, Liver | 211 | 207 | (24, 25, 28, 29) |
| NDBzA | Liver | 80 | 76 | (26) |
| NDMA | Liver | 46 | 30 | (23) |
| Sunlight | Skin | 63 | 62 | (20) |
| UVB | Skin | 116 | 109 | (19, 21) |
| X-Ray | Brain, Liver, Spleen | 98 | 35 | (22) |

**Supplementary Table 2.** Summary of mutant frequencies measured in control and treated animals.

| Group | Number of Animals | Mutants | Plaque Forming Units | Average Mutant Frequency ($x10^{-5}$)[1] | Standard Error | Fold Change | P-value |
|---|---|---|---|---|---|---|---|
| Control[2] | 19 | 403 | 5,953,396 | 5.71 | 0.63 | - | - |
| BaP | 6 | 15,240 | 2,130,473 | 701.70 | 54.02 | 122.9 | **<0.0001** |
| PRC | 8 | 904 | 1,597,855 | 55.13 | 4.51 | 9.7 | **<0.0001** |
| ENU | 6 | 446 | 1,090,253 | 40.93 | 4.34 | 7.2 | **<0.0001** |
| TEM | 7 | 293 | 2,425,691 | 9.18 | 1.86 | 1.6 | **0.048** |

[1]Outliers removed.

[2]The number of control animals and mutants used for mutant frequency calculations are different from the number used for spectral/signature analysis. Mutants from additional control animals were collected for spectral analysis, but plaque forming units were not quantified for these animals.

**Supplementary Table S3.** Summary of mutants, independent mutations, and unique mutational events characterized in each exposure group.

| Group | Mutants | Independent Mutations | Unique SNV Events[1] |
|---|---|---|---|
| Control | 1046 | 512 | 55 |
| BaP | 2914 | 1547 | 377 |
| PRC | 129 | 120 | 14 |
| ENU | 902 | 419 | 85 |
| TEM | 428 | 153 | 22 |

[1]There are 3096 positions × 3 possible substitutions for a total of 9288 possible unique SNV events. SNV events were considered unique to a group if they were only observed in that particular group.

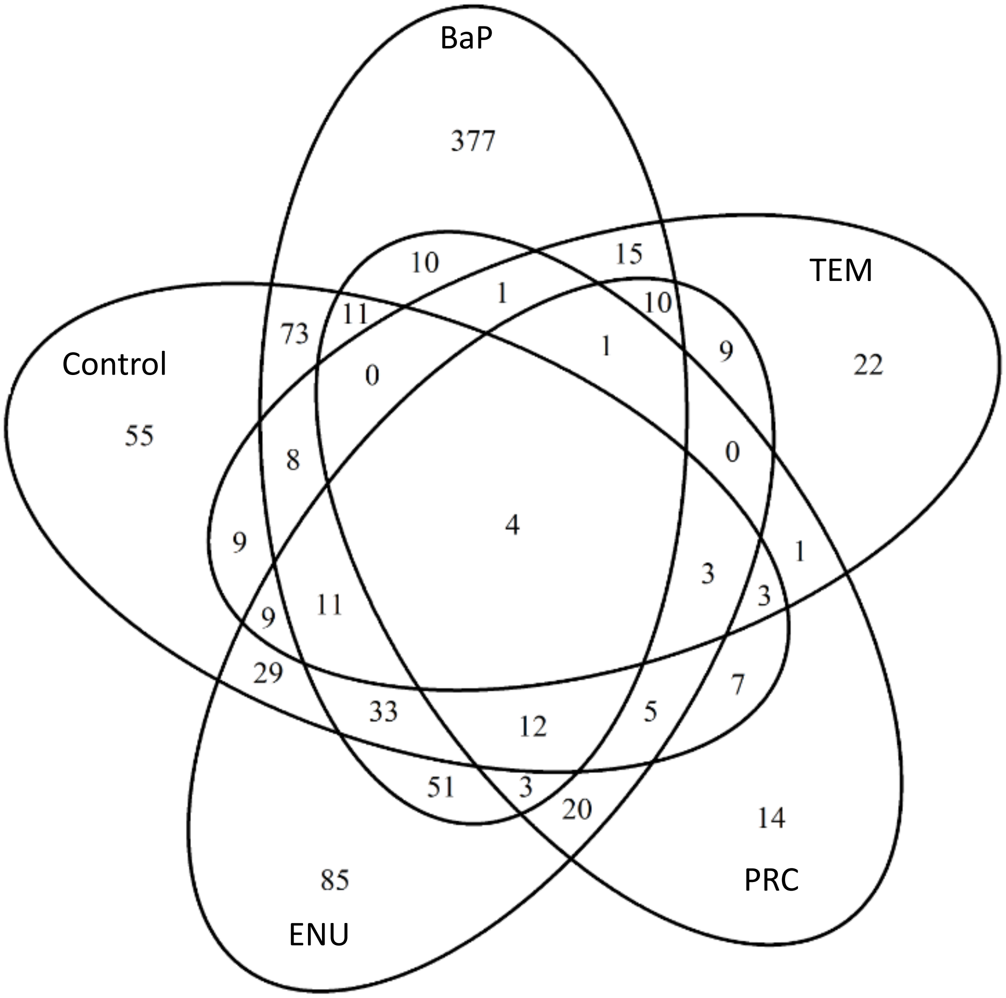


**Supplementary Figure S1.** Venn diagram showing the overlap of unique mutations in the controls and four chemical treatment groups**.**

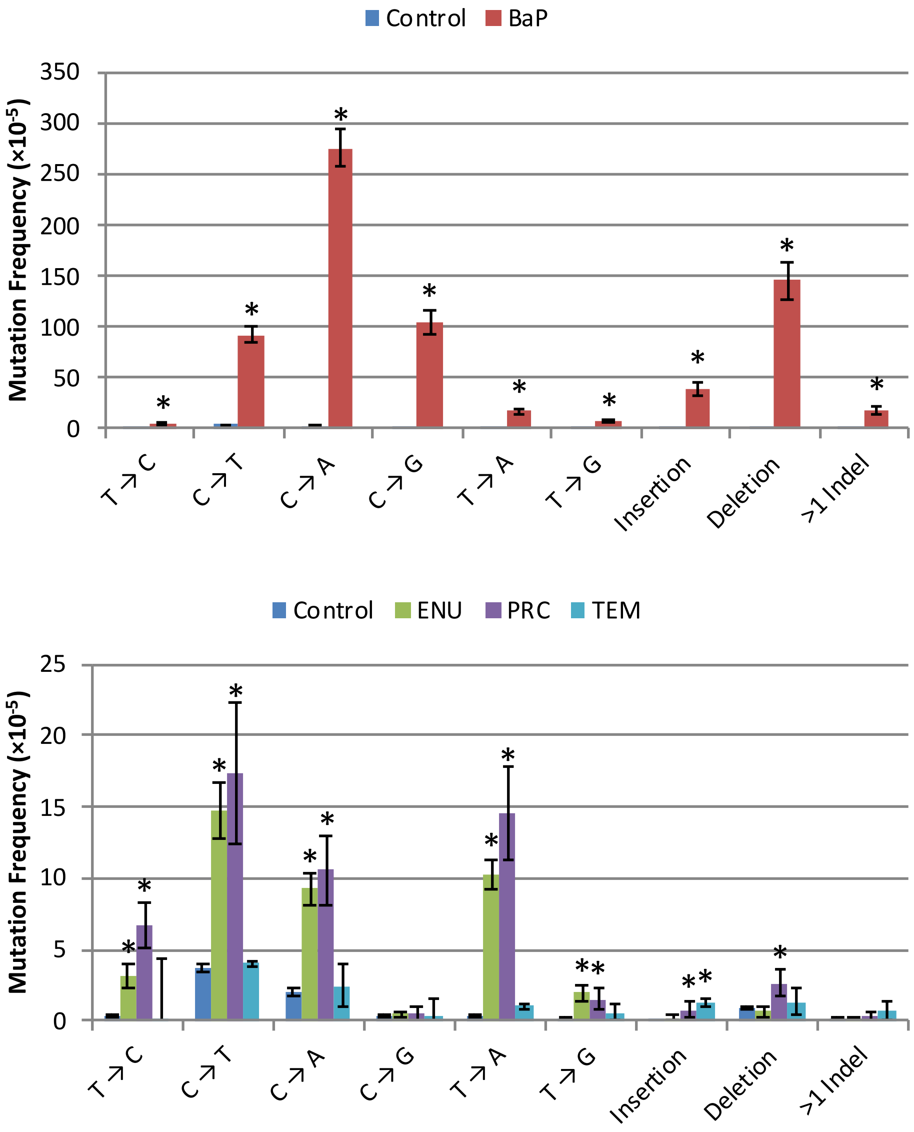


**Supplementary Figure S2. Mutation frequencies of each mutation type compared across exposure groups.** BaP was presented in its own chart because of its high fold-induction compared to control. Asterisks signifies P < 0.05 for mutation type (pairwise comparisons with Bonferonni correction).

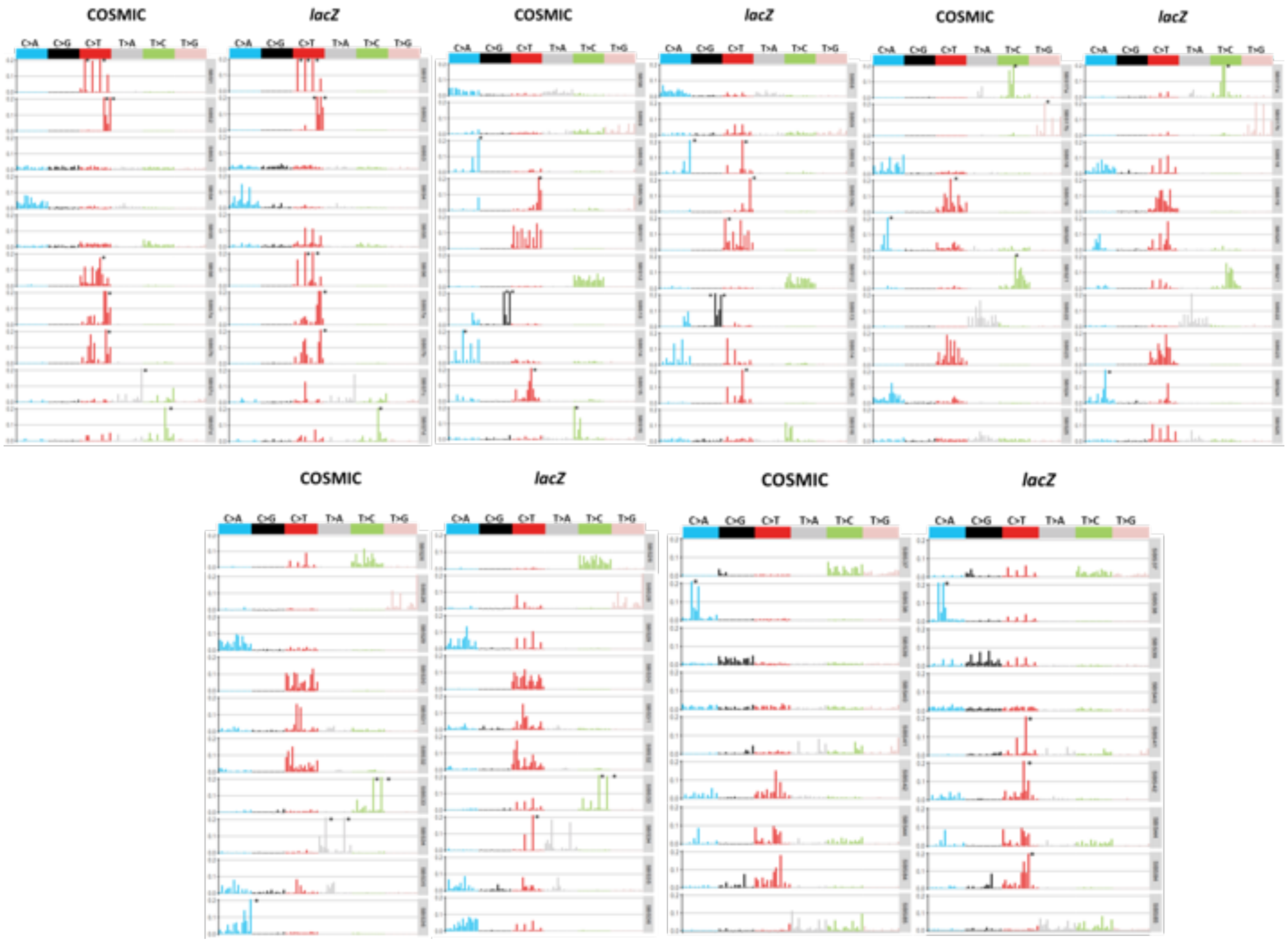


**Supplementary Figure S3 -** Comparison of COSMIC SBS signatures and *lacZ*-adjusted signatures normalized to the ratio of *lacZ* vs. human trinucleotide frequencies. COSMIC signature data were obtained from the COSMIC database (version 3)

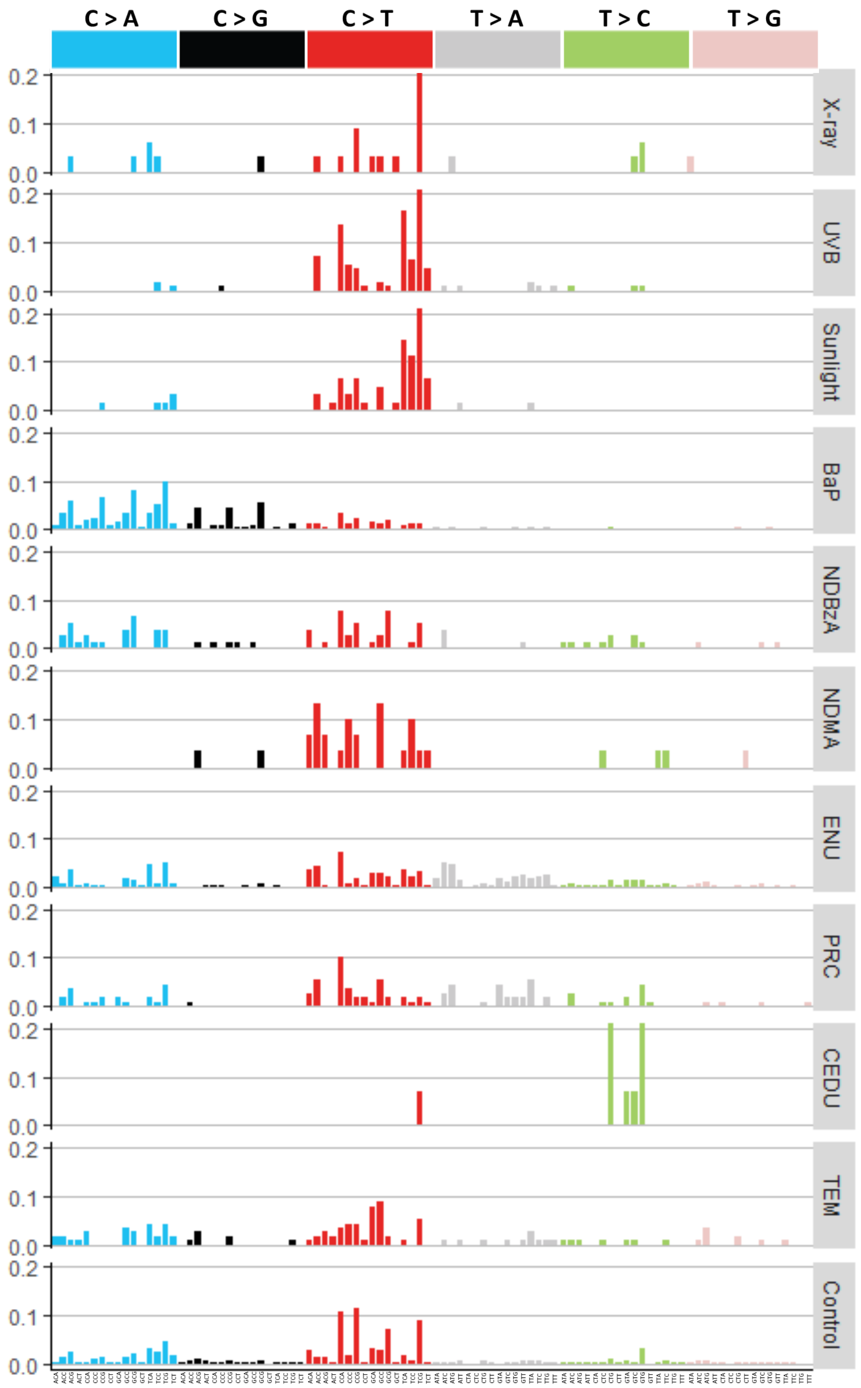


**Supplementary Figure S4.** The 96-base context mutation spectra catalogued from *in vivo* SNVs characterized by NGS and Sanger from control and chemical-exposed animals.

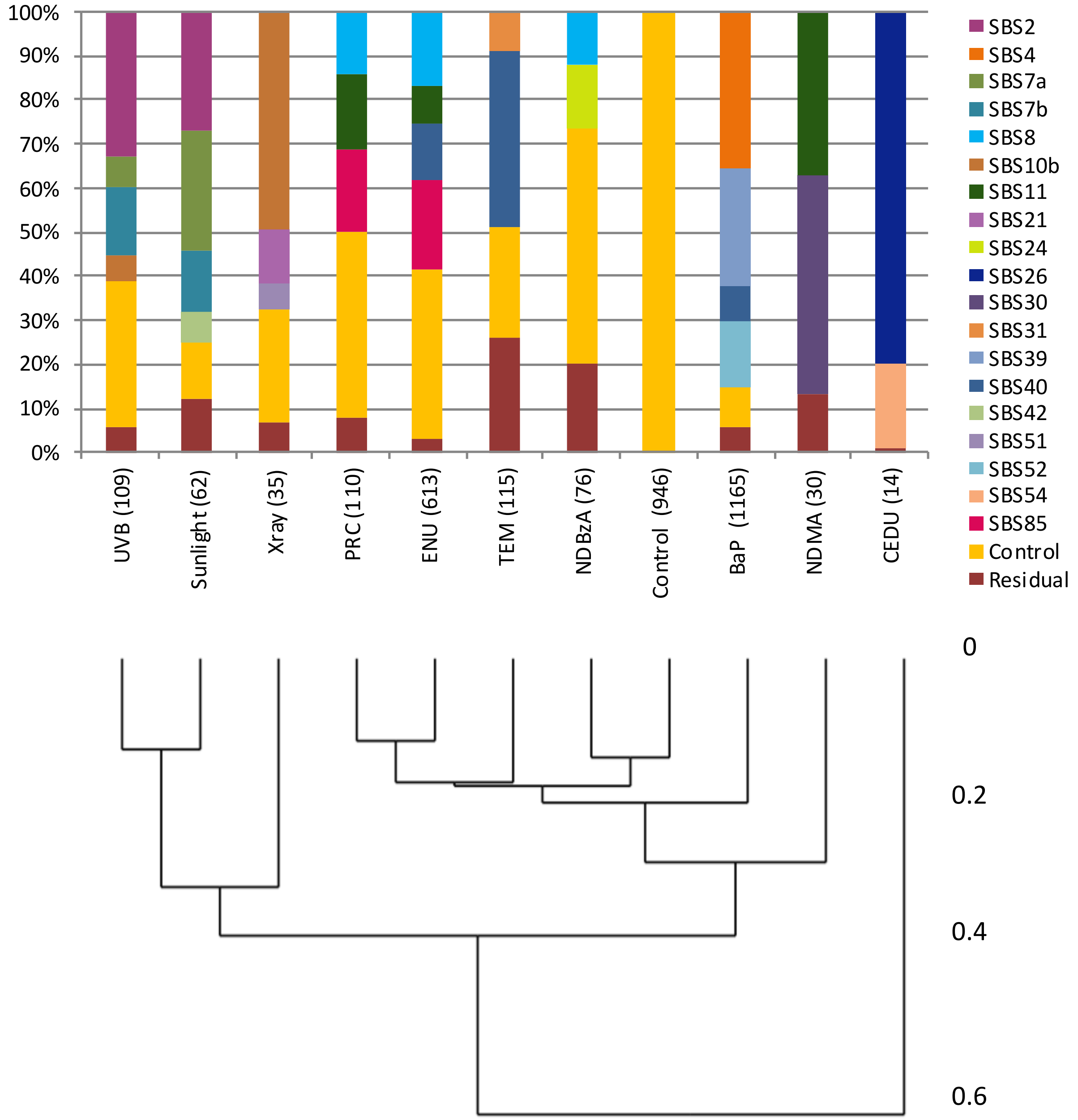


**Supplementary Figure S5. Decomposition of SNV data from different studies using COSMIC signatures.** Number of SNVs sequenced displayed in parentheses. Dendrogram is based on motif matrices.

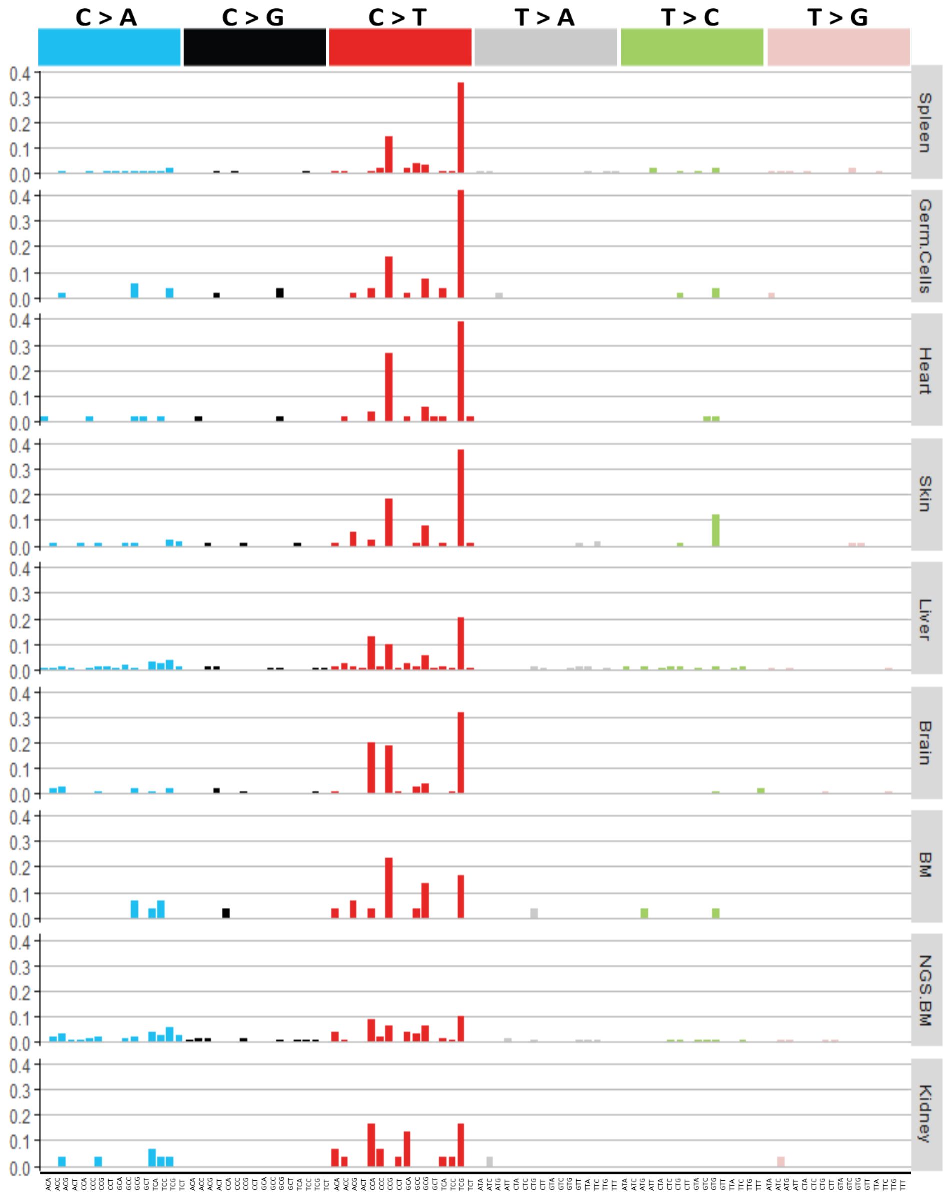


**Supplementary Figure S6. The 96-base context mutation spectra catalogued from *in vivo* SNVs characterized in control tissues.** Note: the Sanger sequencing results show the SNVs from all positions, including the 5 mutation hotspots. Bone marrow is abbreviated to BM.

| Signature | Electromagnetic radiation | | | Bulky adducts | | Alkylating agents | | | Base analog | Clastogen | Spontaneous |
|---|---|---|---|---|---|---|---|---|---|---|---|
| | X rays (35) | UVB (109) | Sunlight (62) | BaP (1165) | NDBzA (76) | NDMA (30) | ENU (613) | PRC (110) | CEDU (14) | TEM (115) | Control (946) |
| **SBS 2** | | 34 | 29 | | | | | | | | |
| **SBS 4** | | | | 34 | | | | | | | |
| **SBS 5** | | | | | | | | | | | 26 |
| **SBS 7a** | | | 23 | | | | | | | | |
| **SBS 7b** | 7 | 15 | 20 | | 9 | | | | | | |
| **SBS 8** | | | | | 8 | | 14 | 9 | | | |
| **SBS 10b** | 50 | 11 | | | | | | | | | |
| **SBS 11** | | | | | | 37 | | | | | |
| **SBS 19** | 9 | | | | | | | | | | 10 |
| **SBS 21** | 15 | | | | | | | | | | |
| **SBS 23** | | | | | | | | | | 7 | |
| **SBS 24** | | | | | 14 | | | | | | |
| **SBS 26** | | | | | | | | | 80 | | |
| **SBS 30** | | 34 | | | | 50 | 21 | 25 | | 8 | 18 |
| **SBS 31** | | | | | | | | | | | |
| **SBS 39** | | | | 28 | | | | | | | |
| **SBS 40** | | | | 13 | 32 | | 31 | 28 | | 40 | |
| **SBS 42** | | | 10 | | | | | 8 | | 16 | |
| **SBS 46** | | | | | 13 | | | | | | |
| **SBS 51** | 15 | | | | | | | | | | |
| **SBS 52** | | | | 16 | | | 10 | | | 7 | 7 |
| **SBS 54** | | | | | | | | | 19 | | |
| **SBS 56** | | | | | | | | | | | 7 |
| **SBS 85** | | 6 | | | | | 14 | 15 | | | |
| **Residual** | 5 | 0 | 19 | 9 | 24 | 13 | 10 | 15 | 1 | 22 | 0 |
| **Pearson coef** | 0.96 | 0.93 | 0.98 | 0.92 | 0.69 | 0.89 | 0.59 | 0.50 | 0.67 | 0.70 | 1.00 |

**Supplementary Figure S7. Relative contributions of COSMIC SBS signatures identified with SNV data generated from different studies using NGS or Sanger sequencing without the use of the control signature.** The number below each agent indicate the number of unique mutant sequenced, while the number in each box represent the percent contribution of each signature to the mutation profile of each tested agent. The last row shows that Pearson's coefficient of reconstructed signatures vs actual mutation data.
